# Warm waters undermine cryptic female choice

**DOI:** 10.1101/2025.04.10.648287

**Authors:** Matthew C. Kustra, Louise M. Alissa, Michaela M. Rogers, Megan M. Molinari, Kelly A. Stiver, Susan Marsh-Rollo, Jennifer K. Hellmann, Suzanne H. Alonzo

**Affiliations:** Department of Integrative Biology, University of California, Berkeley, CA, 94720, USA; Museum of Vertebrate Zoology, University of California, Berkeley, CA, 94720, USA; Miller Institute for Basic Research in Science, University of California, Berkeley, CA, 94720, USA; Department of Ecology and Evolutionary Biology, University of California, Santa Cruz, CA 95064, USA; Department of Evolution, Ecology, and Organismal Biology, the Ohio State University, Columbus, OH, 43210, USA; Psychology Department, Southern Connecticut State University, New Haven, CT 06515, USA; Department of Psychology, Neuroscience and Behaviour, McMaster University, Hamilton, ON L8S 4K1, Canada; Institute of Marine Sciences, University of California, Santa Cruz, California 95064, USA

## Abstract

Reproduction is often more thermally sensitive than survival. Thus, understanding the thermal sensitivity of reproductive interactions is crucial given global warming. However, it is unknown how temperature influences female control over fertilization after mating (*i.e.,* cryptic female choice). We tested how temperatures relevant to current conditions and climate change projections influence cryptic female choice in a marine fish, *Symphodus ocellatus*. Under typical conditions, females bias fertilization dynamics to favor dominant males. We find that warmer temperatures decrease female influence on sperm velocity and reduce the expected paternity of dominant males. Our results demonstrate that temperatures relevant to climate change can shift the balance between mate choice and male-male competition. Thus, climate change may influence sexual selection, leading to evolutionary changes in reproductive traits.

## Main Text

Temperature influences nearly every aspect of organismal biology, from molecular interactions to individual behavior (*1*). Thus, understanding the thermal sensitivity of organismal traits and interactions among individuals is crucial in a changing world where seasonal shifts are occurring and the frequency of extreme climatic events and average temperatures are increasing (*2–4*). Although rising temperatures and heat waves can be lethal and severely affect species viability (*2*, *3*), less extreme temperatures have important sub-lethal effects, especially on traits related to reproduction (*5–8*). There is growing evidence that temperature influences interactions between the sexes, mating behaviors, and mating systems in ways that have cascading effects on population-level processes and species persistence (reviewed in *8, 9*). However, these studies have almost exclusively focused on interactions before mating, even though important interactions between the sexes also occur after mating (*e.g.*, gamete interactions). Thus, we may be underestimating ways in which temperature could change the strength and direction of sexual selection.

We know traits important for fertilization (*e.g*., egg viability, sperm performance) are thermally sensitive (*6*, *10–14*). However, gametes do not function in a vacuum, and female physiology can greatly influence sperm function and fertilization success in a wide range of taxa (*e.g.*, mammals (*15*, *16*), sea urchins (*17*), insects (*18*), and fish (*19*), reviewed in *20,21*). This can be through interactions with the female reproductive tract or with female reproductive fluids (*18*, *19*). Female reproductive fluids consist of a mixture of proteins, chemoattractants, hormones, carbohydrates, ions, and other organic compounds (reviewed in *22–24*), which can positively affect sperm characteristics (reviewed in *22–24*) and influence fertilization dynamics (*24–26*). For example, female reproductive fluids can influence the relative importance of different sperm characteristics (*e.g.*, sperm velocity over sperm number) in determining fertilization success (*25*, *26*). Thus, female reproductive fluids allow females to exhibit female choice after mating (*i.e.,* cryptic female choice). Cryptic female choice can have important population consequences as female reproductive fluids can select for higher quality sperm or sperm that are more genetically compatible (*e.g*., inbreeding avoidance), producing fitter offspring (*17*, *27–31*). Temperature likely influences cryptic female choice because properties important in sperm-female reproductive fluid interactions like pH, osmolality, and viscosity are temperature dependent (*19*, *23*, *32*, *33*). However, it remains unknown how temperature alters the influence of female reproductive fluid on sperm performance, thereby altering female control over fertilization. This limits our understanding of how sexual selection changes with seasonal temperature fluctuations and temperatures outside the range of historical norms due to climate change. Here, we test whether changes in temperature alter these interactions in ways that could change the strength and direction of sexual selection.

Specifically, we investigate how temperature affects sperm velocity and whether temperature alters the effect of female reproductive fluid on sperm velocity in a Mediterranean fish species, the ocellated wrasse (*Symphodus ocellatus*). This is an excellent species for studying how temperature influences postmating female-sperm interactions. First, ovarian fluid positively impacts sperm velocity and changes fertilization dynamics to favor males that females have a strong mating preference for (*26*). Second, like most marine fish, the ocellated wrasse is an ectotherm with external fertilization, meaning that their gametes/fluids are directly exposed to the ambient temperature during reproduction and are likely vulnerable to heatwaves (*33*). Third, there is a large change in temperature over the reproductive season (Fig. 1A; *34*). Fourth, the Mediterranean is both a biodiversity hotspot and a climate change hotspot (*35*), with average temperatures and the intensity and frequency of marine heatwaves steadily increasing (Fig. 1B; *36, 37*). Understanding how temperature influences interactions between ovarian fluid and sperm in this species will provide insight into how sexual selection will change with temperature in this species and many other marine species with similar reproductive biology (*38*).

**Fig. 1.**
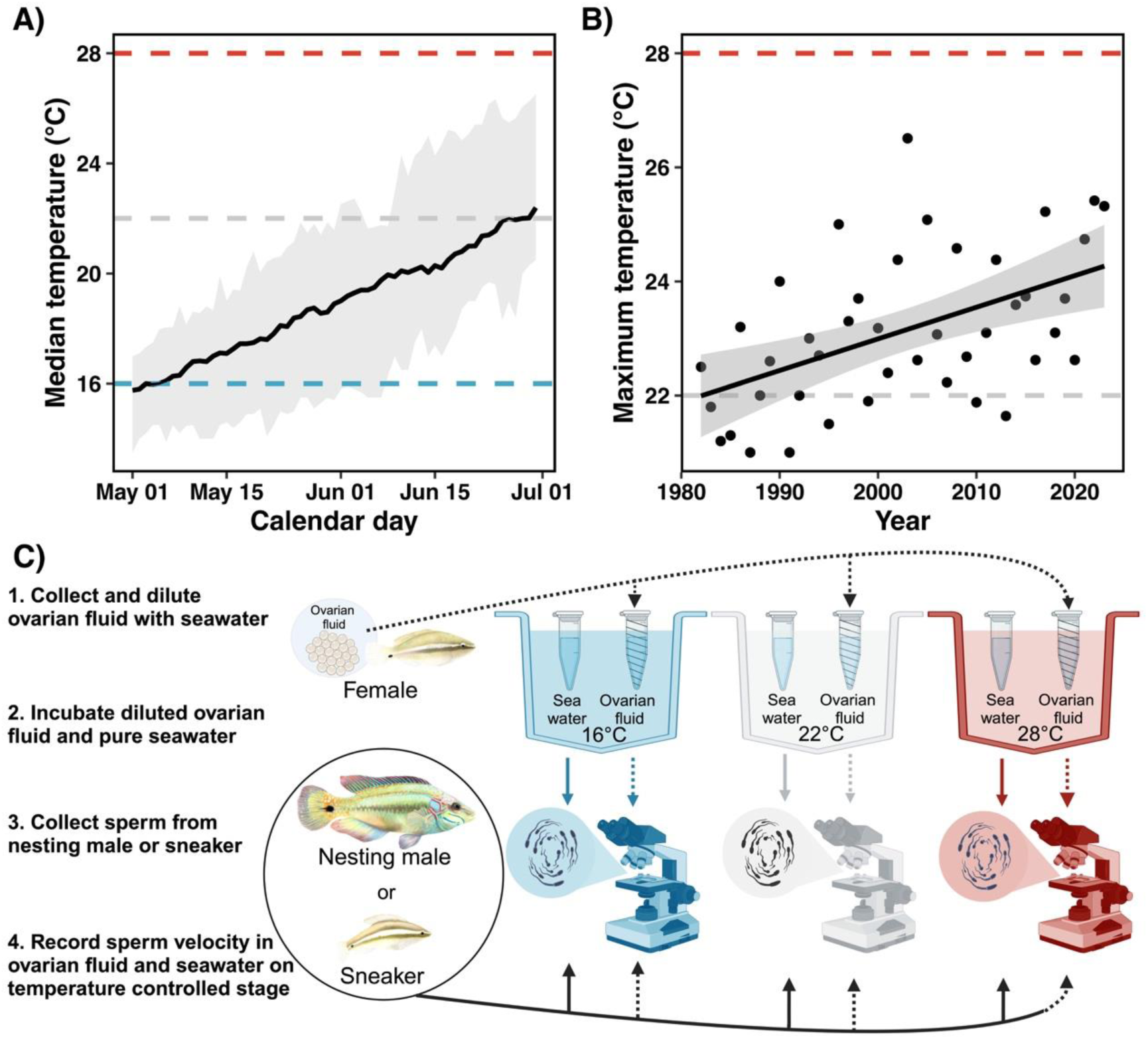
(A) Water temperature increases during the reproductive season, (B) maximum temperature during the reproductive season has increased since 1982, and (C) diagram of the experimental design. **(A)** The black line shows the median water temperature, and the grey shading represents the maximum and minimum temperatures across all years from 1982 to 2023 on different days of the reproductive season. **(B)** Scatter plot with a line of best fit of maximum temperatures during the reproductive season (*β_year_* 0.09, *t* = 5.393, *p <*0.001). Grey shading represents the 95% confidence interval. Colored dashed lines indicate the test temperatures of the experiment, with blue being the coldest test temperature (16℃), grey being the intermediate temperature (22℃), and red being the hottest test temperature (28℃). (**A-B**) These temperature data were collected at the field station where the experiment was conducted (*34*, *46*). (**C**) Each replicate consisted of a single nesting male or single sneaker male. We first collected ovarian fluid from a female and diluted it with filtered seawater into three samples. We incubated these three samples of diluted ovarian fluid and three samples of pure seawater at three different temperatures. We then collected sperm from a male and placed the sperm on a temperature-controlled stage, matching the incubation temperature. We then activated sperm with each fluid treatment at each test temperature and recorded sperm velocity. This resulted in six separate sperm velocity measurements per male and female combination (n= 38; 19 nesting males, 19 sneakers). However, due to exclusion factors, we were not able to use all six treatments for certain analyses for each replicate. Tables S1 and S2 show the exact sample sizes per treatment for each analysis. (**C**) We made the diagram in BioRender. Fish illustrations were made by Clara Lacy adapted from (*47*).

The ocellated wrasse’s mating system, like many fish, involves female choice among male territories. Three distinct male phenotypes exist: a large dominant nesting male, an intermediate-sized satellite male, and a small sneaker male (*39–41*). Nesting males represent a distinct life history pathway from sneakers that indicates nesting males are of higher quality (*42*). Further, females have strong behavioral mating preferences for nesting males, which build nests and provide all parental care (*43*). To override female choice, sneaker males sneak spawn after the nesting male (*44*) and release ∼3x more sperm than nesting males (*45*). Female reproductive fluid (ovarian fluid) counteracts this numerical advantage by decreasing the importance of sperm number but increasing the importance of sperm velocity in determining fertilization success, effectively favoring the preferred nesting males (*26*). These discrete male phenotypes make it easier to assess how temperature influences cryptic female choice compared to a system with more continuous variation in male phenotypes and female preferences. If temperature affects ovarian fluid, we can test the general question: does temperature change a female’s ability to control fertilization?

We performed a full factorial experiment to test the effect of temperature on postmating female-male interactions on both nesting and sneaker males. We focused on these male types because they represent distinct life history pathways and different degrees of quality. We measured each male’s sperm velocity in the presence or absence of ovarian fluid at three temperatures (16℃, 22℃, and 28℃; Fig. 1). These three temperatures represent temperatures the fish currently experience during the reproductive season and realistic future conditions given the climate change projections of average seawater temperature increases of 1.8℃ to 3.5℃ by 2100 (Fig. 1; *36, 37*). This resulted in six different treatment combinations per individual male (Fig.1). We looked at both sperm velocity at 30 seconds, since we know this to be important in determining fertilization success, and sperm velocity at five minutes to assess sperm longevity. We then assessed if there was variation in individual responses to temperature because this provides insight into the population’s evolutionary potential and how temperature may modulate the opportunity for sexual selection. Finally, we used a model based on previously published data (*26*) to determine how changes in initial velocity are predicted to influence the relative paternity of nesting males. We found that nesting males have slower sperm at warmer temperatures in ovarian fluid, decreasing expected paternity. This lower nesting male paternity could lead to a shift in their investment in parental care and/or an increase in the nest abandonment rate. As nesting male care is necessary for offspring survival and nesting males are likely of higher quality (*42*), warming temperatures could negatively affect this species. More generally, we demonstrate that temperature can alter female control over fertilization and the dynamics of sexual selection, with important consequences for species persistence and resilience.

### Increasing temperature reduces the positive effect of ovarian fluid for preferred dominant nesting males resulting in sneaker males having faster sperm

We first examined how ovarian fluid, male type (nesting male or sneaker), and temperature influence initial sperm velocity (velocity at 30 seconds). Initial sperm velocity is a good predictor of fertilization in this species (*26*) and other fish (*48*). We found that initial sperm velocity was significantly influenced by the three-way interaction between activating fluid, experimental temperature, and male type (Table 1A). Specifically, nesting male sperm were slower than sneaker male sperm at warmer temperatures when in ovarian fluid (Fig. 2A, B).

**Fig. 2.**
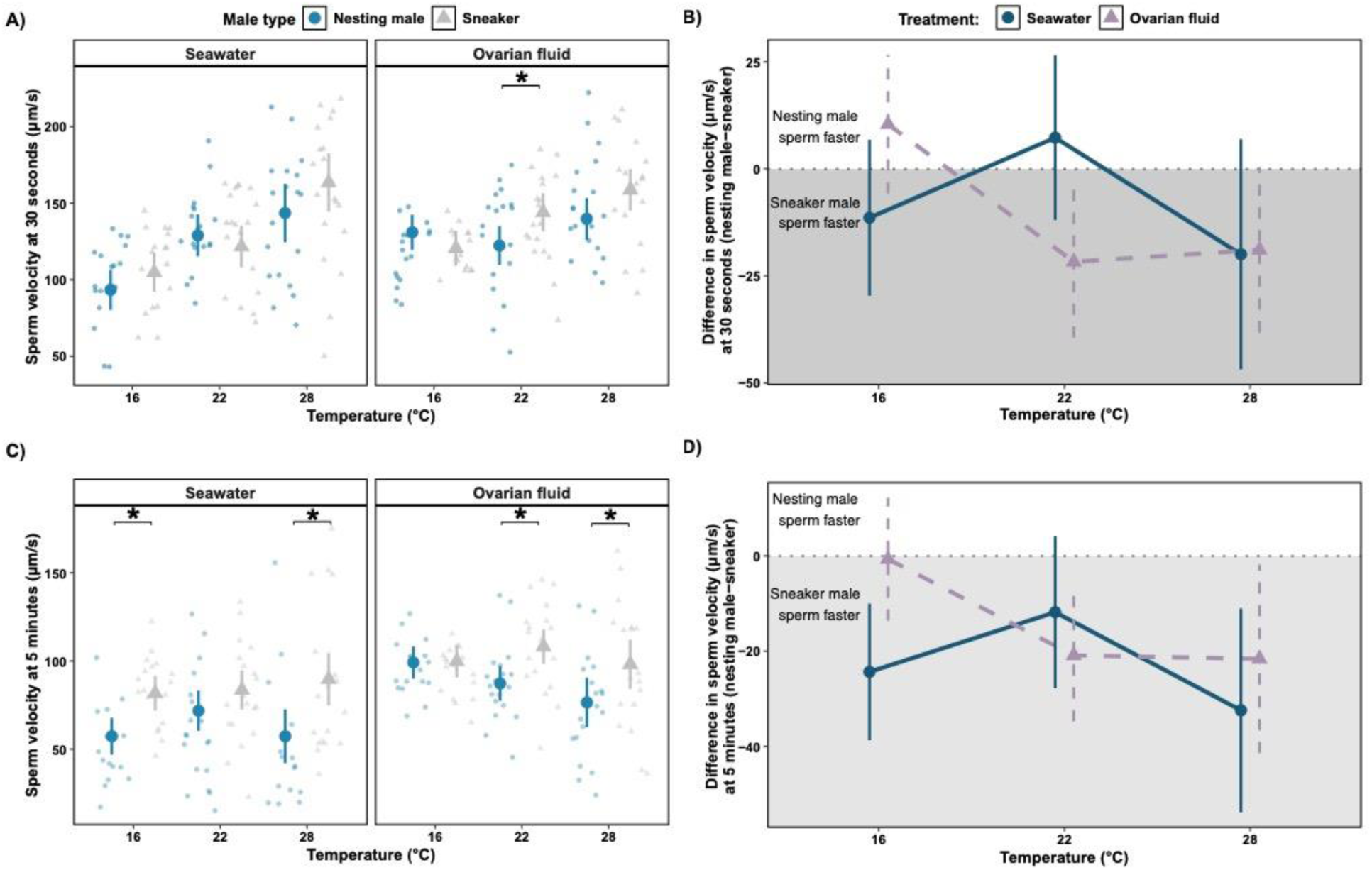
Increasing temperature increases sperm velocity but limits the positive effect of ovarian fluid, resulting in sneakers having higher sperm velocity than nesting males. **(A)** Solid shapes indicate the predicted sperm velocity at 30 seconds of each treatment based on the model described in Table 1A, and bars represent the 95% confidence intervals. The smaller points in the background are averages of sperm velocity of all motile sperm at 30 seconds post activation per individual and are jittered along the x-axis. **(B)** The same data from panel A is presented as a difference score with solid shapes indicating the contrast between sneaker males and nesting males at each treatment based on the model described in Table 1A. The bars represent the 95% confidence intervals of those contrasts. **(C)** Solid shapes indicate the predicted sperm velocity at five minutes of each treatment based on the model described in Table 1B, and bars represent the 95% confidence intervals. The smaller points in the background are averages of sperm velocity of all motile sperm at five minutes post-activation per individual and are jittered along the x-axis. **(D)** The same data as panel C is presented as a difference score with solid shapes indicating the contrast between sneaker males and nesting males at each treatment based on the model described in Table 1B, and bars represent the 95% confidence intervals of those contrasts. **(A and C)** Asterixis indicate significant differences between male types. The statistical significance of all pairwise comparisons is given in tables S3-S8. **(B and D)** The dotted grey line at zero means no difference between nesting males and sneakers. Positive values (white background) indicate nesting males have faster sperm than sneaker males, and negative values (grey background) indicate sneakers have faster sperm than nesting males.

**Table 1.** Results of the linear mixed-effects model explaining sperm velocity at (A) 30 seconds and (B) five minutes. The significance of fixed effects was determined with a Type II ANOVA Wald χ^2^ test. Random effects for both models were a random intercept for the field of view nested within the trial and random slopes for both activating fluid treatment and temperature treatment of each trial. Significant effects are bolded.

| **Trait** | **Model Effect** | **$\chi^2$** | **P(>$\chi^2$)** |
| --- | --- | --- | --- |
| (A) Sperm velocity at 30 seconds | **Temperature** | **40.711** | **<0.0001** |
|  | Activating fluid | 3.712 | 0.0540 |
|  | Male type | 1.477 | 0.224 |
|  | **Activating fluid: temperature** | **60.949** | **<0.0001** |
|  | Activating fluid: male type | 0.0977 | 0.7546 |
|  | Temperature: male type | 3.599 | 0.1654 |
|  | **Activating fluid: temperature: male type** | **43.181** | **<0.0001** |
| (B) Sperm velocity at five minutes | Temperature | 3.358 | 0.1866 |
|  | **Activating fluid** | **24.646** | **<0.0001** |
|  | **Male type** | **7.481** | **0.0062** |
|  | **Activating fluid: temperature** | **53.100** | **<0.0001** |
|  | Activating fluid: male type | 0.8455 | 0.3578 |
|  | Temperature: male type | 4.868 | 0.0877 |
|  | **Activating fluid: temperature: male type** | **74.412** | **<0.0001** |

When directly comparing the initial sperm velocity of nesting males and sneakers in ovarian fluid, there was no significant difference between nesting males and sneakers at 16℃ (Fig. 2A, B). However, at 22℃ and 28℃, sneakers had faster sperm than nesting males, although this was not significant at 28℃ after correcting for multiple testing (Fig. 2A, B). In seawater, there was no difference between the male types at any temperature (Fig. 2A, B). These differences between the males were driven by differing responses to temperature and activating fluid. Initial sperm velocity increased with temperature for both males in seawater, but in ovarian fluid, velocity only increased with temperature for sneaker males (Fig. 2A; Table S2; Table S3). At 16℃, ovarian fluid positively affected nesting male swimming velocity relative to seawater. At 22℃ and 28℃, there was no significant difference in velocity between sperm activated in seawater and ovarian fluid for nesting males. At 16℃ and 22℃, ovarian fluid had a significant positive effect on sneaker male velocity. However, at 28℃, ovarian fluid had no significant influence on sperm velocity of sneaker males. We report all pairwise comparisons in Tables S3-S5. In summary, sneaker male sperm swam faster than nesting male sperm at warmer temperatures in ovarian fluid.

Initial sperm velocity may tradeoff with sperm longevity, so we next looked at sperm velocity at five minutes. We found that sperm velocity at five minutes was also significantly influenced by the three-way interaction between activating fluid, experimental temperature, and male type (Table 1B; Fig. 2C). Specifically, nesting male sperm swam slower than sneaker male sperm at warmer temperatures when in ovarian fluid (Fig. 2C, D).

When directly comparing the sperm velocity at five minutes of nesting males and sneakers in ovarian fluid at 16℃, sneaker males and nesting males had similar sperm swimming velocity (Fig. 2C, D). However, at 22℃ and 28℃, sneakers had significantly faster sperm at five minutes (Fig. 2C, D). In seawater, nesting males had lower sperm velocity than sneakers at all temperatures, although this was non-significant at 22℃ (Fig. 2C, D). These differences between the males were driven by differing responses to temperature and activating fluid. Nesting male sperm velocity at five minutes decreased with increasing temperature in ovarian fluid (Fig. 2C) but not seawater. This was likely because sperm velocity was low in seawater across all temperature treatments for all males, which was expected because ovarian fluid increases sperm longevity in this species (20). Temperature did not influence sperm velocity at five minutes for sneaker males in either seawater or ovarian fluid (Fig. 2C). Ovarian fluid increased sperm velocity for nesting males at five minutes compared to seawater across all temperatures, though the magnitude of this effect was strongest at 16℃ (Fig. 2C). Ovarian fluid only increased sperm velocity at five minutes for sneaker males at 16℃ and 22℃. We report all possible pairwise comparisons in tables S6-S8.

To account for the potential of temperature shock, we ran supplemental analyses including the difference in temperature between treatment temperature and median ocean temperature on the day of the replicate. We found that including this effect did not change the qualitative results at 30 seconds or five minutes post-activation (Table S9) or improve model fit (Table S10).

Sperm velocity is an important factor determining fertilization success in many species. Our finding of initial sperm velocity increasing with temperature is consistent with other studies demonstrating a positive impact of temperature on initial sperm velocity in fish (*11*). This could be due to increased sperm metabolism, resulting in reduced longevity. Indeed, we found this to be the case for nesting males whose sperm velocity at five minutes decreased with increasing temperature in ovarian fluid. Interestingly, sperm velocity at five minutes did not decrease with increasing temperature for sneaker males. One potential reason for this difference could be physiological differences in the sperm. Sneakers in this system have longer-lived sperm than nesting males in ambient temperatures (*26*). Although this has not been tested in this species, this could indicate higher energy stores in sneaker male sperm cells (i.e., ATP). A recent meta-analysis (*49*) and review (*50*) found that sneaker males in fish often have sperm with higher ATP content.

Ovarian fluid positively influences sperm velocity and sperm longevity for fish generally (*23*) and in this species (*26*). Here, we found that ovarian fluid only increased initial sperm velocity at 16℃ for both males. Ovarian fluid improved sperm longevity for nesting males across all temperatures, but this effect weakened as temperature increased to the point that sperm longevity was significantly lower than that of sneakers at 22℃ and 28℃. One possibility is that this effect of temperature could be due to temperature influencing the viscosity or osmolality of the ovarian fluid, characteristics that influence sperm velocity in other taxa (*19*, *23*, *32*).

In summary, higher temperatures limited the ability of females to influence sperm velocity, likely making male-male competition more important than female choice at warmer temperatures. To our knowledge, the only other study to explore temperature effects on female reproductive fluid was in an internally fertilizing lizard (*51*), which found that increasing temperature did not influence the impact of female reproductive fluid on sperm velocity. More research on how temperature influences sperm-female reproductive fluid interactions is needed to understand and predict the consequences of climate change.

### Significant variation exists in individual responses to temperature and ovarian fluid

Because the evolutionary potential of a population to respond to environmental change depends on existing variation, it is important to also examine individual variation in responses to temperature. To determine whether there was significant variation in individual responses to either temperature or ovarian fluid, we tested the significance of random slopes. We found that including a random slope for both the activating fluid treatment and temperature treatment significantly improved the model fit for both sperm velocity at 30 seconds (χ^2^=1304.9, *p*<0.0001; Fig. S1; Table S11A) and sperm velocity at five minutes (χ^2^=3064.1, *p*<0.0001; Fig. S2; Table S11B). This indicates significant variation in individual male responses to both treatments for sperm velocity at 30 seconds (Fig. S1) and sperm velocity at five minutes (Fig. S2).

Another potential factor that may influence the evolutionary potential of a population is cryptic genetic variation in the form of hidden reaction norms (*52*, *53*). In other words, existing variation is only apparent in novel environments (*e.g.,* temperatures outside the historical range). To test this, we examined whether sperm velocity had greater variability at warmer temperatures. Specifically, we generated a posterior distribution of differences in the standard deviation of random slope terms at different temperatures. We found that regardless of activating fluid treatment and male type, there was greater individual variation in sperm velocity at the warmest temperature treatment at both 30 seconds and five minutes (Fig. 3, S3). These results are consistent with the potential for cryptic genetic variation. If the variation in responses to temperature observed is heritable, this would indicate the potential for evolutionary change in these traits in response to changing environments. For example, selection could favor females who produce ovarian fluid that is more temperature resistant or males whose sperm are more temperature resistant. Further, the increase in variation of sperm velocity at warmer temperatures also indicates that the opportunity for sexual selection is stronger at warmer temperatures.

**Fig. 3.**
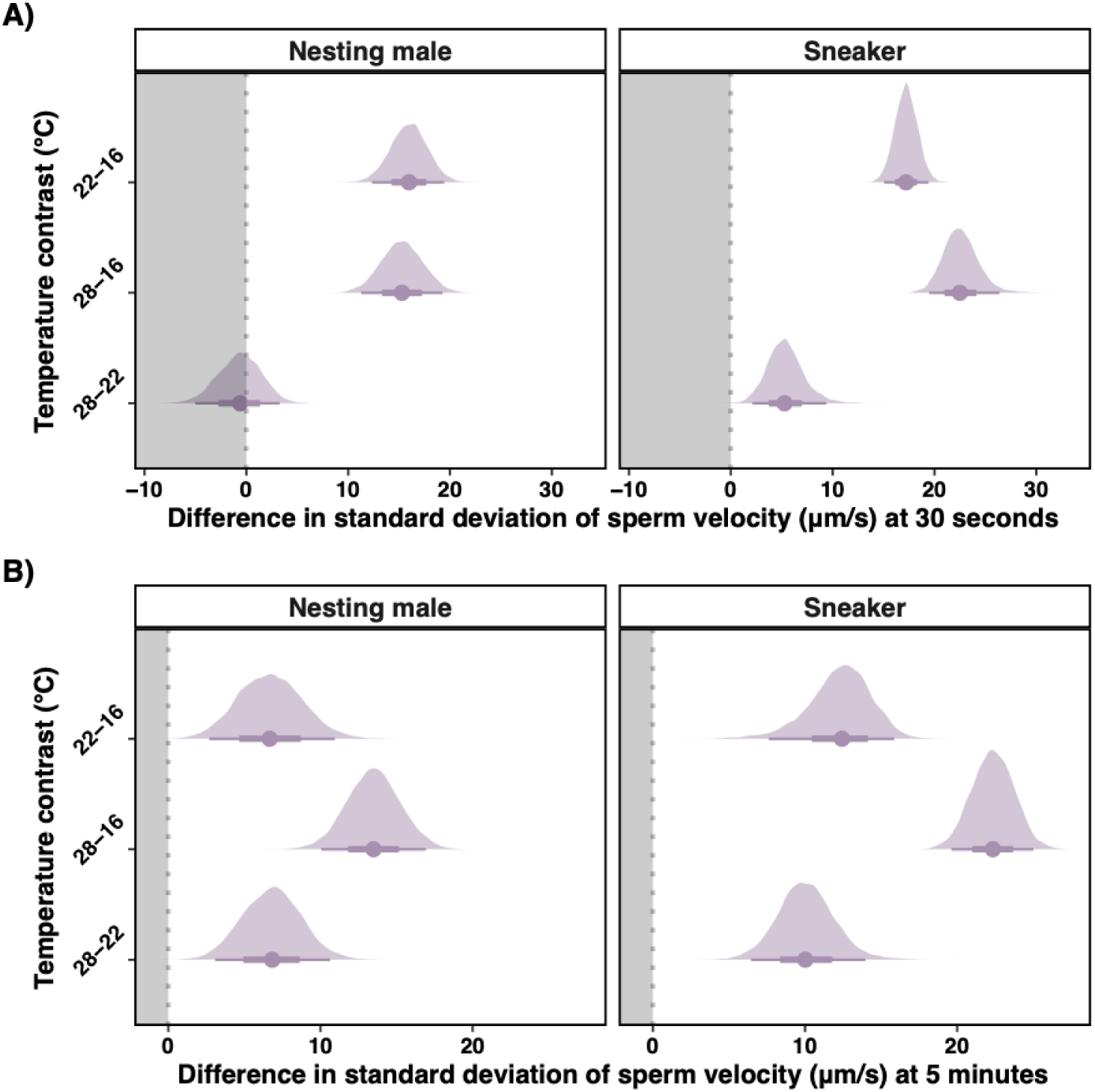
Variation in sperm velocity increases with temperature in ovarian fluid regardless of male type. Posterior probability distributions of the difference in random slope standard deviation at different temperatures for nesting males (left) and sneakers (right) in ovarian fluid at **(A)** 30 seconds and **(B)** five minutes. The posterior distributions were generated with 10,000 simulated draws. Solid points represent the median of the posterior, thin bars represent the 95% credible intervals of the posterior distribution, and thick bars represent the 66% credible interval. The dotted grey lines at zero mean no difference in standard deviation. Negative values (grey background) indicate higher temperature decreased variation. Positive values (white background) indicate higher temperatures increased variation. Similar plots for the seawater control are shown in Fig. S3.

### Increasing temperature undermines cryptic female choice and reduces the reproductive success of preferred dominant nesting males

To determine how temperature and activating fluid influence the relative paternity of preferred nesting males, we refit data from (*26*), which looked at how fertilization success of nesting males was determined by sperm number and sperm velocity in either ovarian fluid or seawater. We accounted for error in our sperm velocity estimates at different temperatures and activating fluids and error in the paternity statistical model by using a sampling approach to generate a posterior distribution. In general, ovarian fluid increased predicted nesting male paternity (Fig. S4). However, this paternity bias towards the nesting male diminished at higher temperatures (Fig. 4). This resulted in a 7.6% (95% CI: 16.5 - 1.5%) reduction in nesting male paternity at 22℃ and a 6.5% (14.9 - 1.2%) reduction at 28℃ compared to 16℃ (Fig. 4). There was no further reduction in nesting male paternity going from 22℃ to 28℃ (Fig. 4).

**Fig. 4.**
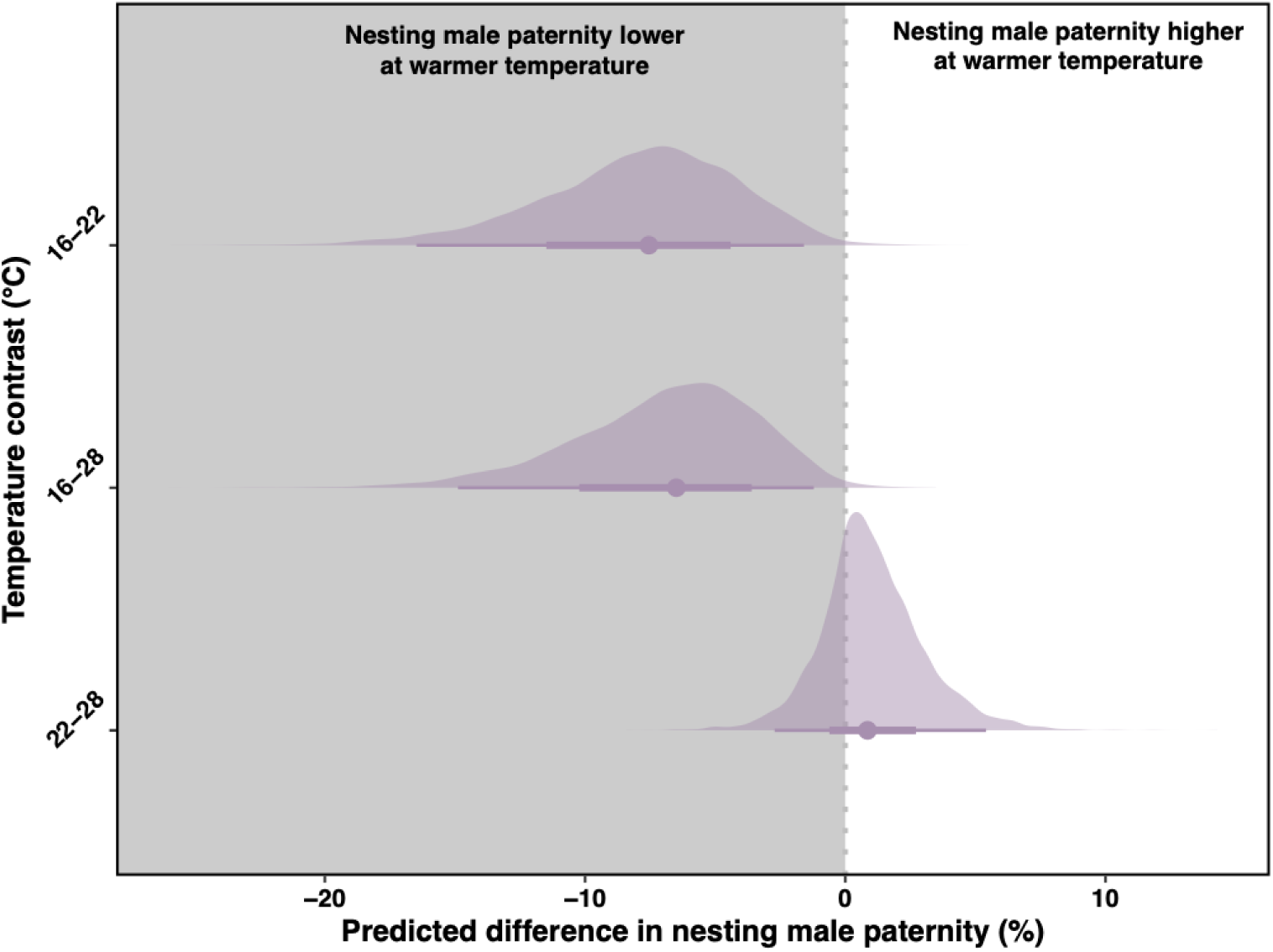
Ovarian fluid has a less positive impact on nesting male paternity at 22℃ and 28℃ compared to 16℃. Shown are the posterior probability distributions of the difference in nesting male paternity at different temperatures in ovarian fluid. Solid points represent the median of the posterior, thin bars represent the 95% credible intervals of the posterior distribution, and thick bars represent the 66% credible interval. The posterior distributions were generated with 10,000 simulated draws. The dotted grey lines at zero mean no difference in nesting male paternity. Negative values (grey background) indicate higher temperatures decreased predicted nesting male paternity. Positive values (white background) indicate higher temperatures increased predicted nesting male paternity. Posterior probability distributions at the different temperatures and activating fluid are shown in Fig. S4.

These results, in combination with previous work in this system, suggest that mechanisms of cryptic female choice that bias paternity to nesting males may currently only function early in the reproductive season. Because temperature typically increases from 16℃ to 22℃ over the reproductive season (Fig. 1A), nesting males may have higher fertilization success in competitive matings at the beginning of the season than at the end. This could influence nesting males’ behavior. For example, nesting males may be under selection to mate guard (e.g., be more aggressive to subordinate males) to prevent sperm competition toward the end of the reproductive season when waters are warmer (*45*). Our results demonstrate a novel way for temperature to change the dynamics of sexual selection within the reproductive season (*54*).

In the context of climate change, these results shed light on how seasonal and temperature shifts could influence this system. Warmer temperatures could result in a change in the frequency of these male phenotypes. With lower expected paternity, nesting males may decrease parental care investment and/or increase nest abandonment rate, which would have negative population effects through decreased offspring survival. Alternatively, nesting males may start building earlier (when temperatures are colder), mirroring phenological shifts of reproduction seen in terrestrial systems (*55*), or build nests deeper where temperatures are colder. However, phenological shifts may be constrained due to other species of the same genus that nest in the same sites immediately preceding the *S. ocellatus* reproductive season (*41*). Male-only parental care is common in fish (*38*). Thus, the dynamics described above may be more general and highlight the importance of understanding the relationship between climate change, sexual selection, and parental care.

## Conclusion

Despite female reproductive fluid playing an important role in reproduction, we know little about how temperature influences female reproductive fluid-sperm interactions. If temperature affects these interactions, this can lead to changes in both absolute and relative fertilization success with important ecological and evolutionary consequences. We show that warm temperatures undermine female control over fertilization dynamics. Interestingly, these effects were present at 22℃, the highest temperature they currently experience during the end of the reproductive season. Thus, temperature is an underappreciated factor modulating postmating sexual selection and the effectiveness of female reproductive fluid both under current conditions and under future climate change projections. Female reproductive fluid in many systems can select for higher quality sperm or sperm that are more genetically compatible, producing fitter offspring (*17*, *27–31*). Thus, by decreasing the effectiveness of female reproductive fluid, rising temperatures could negatively impact population viability. Additionally, by reducing female control over fertilization, warmer temperatures shift the balance between mate choice and male-male competition. Future research into the consequences of climate change on female-male postmating interactions will be essential for understanding how climate change will influence fertility, population persistence, and sexual selection.

## Acknowledgments

We thank the staff at STARESO for assistance during fieldwork, especially C. Steibel. We thank L. Fullgrabe for sharing the long-term temperature data set. Long-term acquisition of temperature data was supported by the STARESO marine station, the Rhone-Mediterranean and Corsica Water Agency and the Collectivity of Corsica under the STARECAPMED project. We thank D. Weiler, J. Fitzpatrick, B. Lyon, M. Pinsky, M. Gamble, C. Martin, D. Tian, C. Tralka, F. Palominos, and R. Abrams for helpful feedback that greatly improved this manuscript.

## Author Contributions

M.C.K. and S.H.A. conceived and designed the experimental design with input and feedback from all authors. M.C.K., M.M.R., L.M.A., M.M.M, and S.H.A. conducted the experiment. M.C.K., L.M.A., M.M.R., M.M.M., K.A.S, S.M.R., and S.H.A. collected fish used in this experiment. M.C.K. analyzed the data with input from J.K.H., K.A.S., and S.H.A. M.C.K. wrote the initial manuscript. All authors edited and approved the final manuscript.

## Data and materials availability

Data and code associated with the manuscript will be made publicly available upon publication and archived on Dryad.

## Supplemental Materials and Methods

### Animal collection

We collected all fish from the wild at the University of Liege Marine Station (STARESO) near Calvi, Corsica, France (42.5806° N, 8.7243° E) from mid-May to mid-June in 2023. This time period is at the peak of the ocellated wrasse breeding season, which typically runs from May to July (*41*). Individuals were kept in a holding tank with flow-through water from the ocean until used in the experiments. The temperatures experienced by the fish were similar to that of the ocean water from which they were collected. Fish were used in the experiment within a day of collection. We did not repeat the use of fish across experimental replicates.

### Experimental design

To test the effect of temperature on sperm motility, we used three different temperature treatments: 16℃, 22℃, and 28℃. We chose these temperatures because they cover both the low, average, and extreme heat waves experienced by fish in this area of the Mediterranean during the reproductive season (Fig. 1). To test how temperature interacts with ovarian fluid, we tested each male’s sperm performance in the presence or absence of ovarian fluid at each temperature. This resulted in six different treatment combinations per individual male. All six treatment combinations were performed within two hours of one another (Fig. 1B).

We first stripped a female of her eggs into a petri dish, pipetted 2 μL of the ovarian fluid surrounding the eggs, and mixed it into a microcentrifuge tube with 4 μL of filtered seawater (similar concentrations to previous work; *26*). This was done three times for each female (one per temperature treatment). We also prepared three microcentrifuge tubes (one per temperature treatment) with 6 μL of filtered seawater. We incubated these tubes in three separate water baths, each set at one of the three different water temperature treatments. The temperature of each water bath was kept within 0.1℃ of the target temperature using INKBIRD ITC-306T thermostats and DaToo flat thermostatic heaters. We incubated these microcentrifuge tubes (containing either seawater or seawater plus ovarian fluid) for at least five minutes prior to use in the experiment.

Next, we collected 0.2μL of milt (semen) from a male fish and added this milt directly to a 3 μL four-chamber LeJa slide. We then set a microscope with a temperature-controlled stage (LinkamWCP) to the desired test temperature to prevent temperature shock. Next, we flushed the milt into the chamber slide with either 2.8 μL diluted ovarian fluid or seawater incubated at the same test temperature. We then recorded sperm motility at 50 frames per second with a Nikon Ci-L microscope at 100x with phase contrast. Sperm characteristics were analyzed with Microptic CASA software immediately after collection. We took continuous frame captures for two minutes, moving the field of view every two captures to record the velocity of many unique sperm. We then haphazardly moved the field of view around and took two captures at three, four, and five minutes to capture sperm longevity. We purposely excluded sperm that were clumped (preventing sperm movement) or sperm that failed to activate. This process was done manually with the person excluding these data blinded to the experimental treatments and male identity.

We additionally excluded any sperm that were not tracked for at least 20 frames. For each male, we repeated the sperm motility analysis for every temperature with both diluted ovarian fluid/seawater treatment (six treatments/male). We used freshly collected milt for each treatment. We purposefully changed the order of treatments across replicates (*i.e.,* different males and females). We ended up with 38 full replicates (19 nesting males, 19 sneakers). However, due to chance events (*e.g*., software crashed during the middle of the experiment), for certain analyses, we did not have all six treatments for each replicate. The exact sample sizes per treatment for each analysis are given in Tables S1, S2.

### Statistical analysis of sperm velocity

We first examined how ovarian fluid and temperature interact to influence the initial sperm velocity of the different male types. We focused on initial sperm velocity because this is a good predictor of fertilization in this species (*26*) and other externally fertilizing fish (*48*).

Specifically, we looked at sperm velocity at 30 seconds following activation, which we refer to as initial sperm velocity. We chose 30 seconds because this was the shortest time in which we could reliably measure a reasonable number of sperm across all six treatments per male: 225±15.2 (mean ± s.e.) sperm. We had measurements across all six treatments for 17 nesting male replicates and 17 sneaker replicates. We had measurements for four or five treatments for the other replicates. We included the incomplete replicates because mixed-effects models account for incomplete blocks, and the data can be used to estimate the treatment effects they contained (see Table S1 for a summary of sample size by treatment). For all sperm velocity analyses, we used the curvilinear sperm velocity (VCL; μm/s) because most sperm did not have linear paths.

To test how activating fluid, male type, and temperature influence initial sperm velocity, we performed a linear mixed effects model in R V.4.1.2 (*56*) using the package *lme4* (*57*). The fixed effects were the three-way interaction between male type (nesting male or sneaker), activating fluid (ovarian fluid or seawater), and temperature (16℃, 22℃, and 28℃), as well as all lower-level effects. We included a random intercept of the field of view nested within the trial. This allowed us to account for the nonindependence of sperm measurements in the same field of view and belonging to the same male. We also included random slopes for activating fluid treatment and temperature treatment effects. The *p-*values of fixed effects were calculated with a Type II ANOVA *χ^2^* test using the *car* package (*58*). Finally, if there was a significant interaction, we performed post hoc comparisons using the *emmeans* package, where *p* values were adjusted with the Tukey method (*59*). Model fit was assessed using the *DHARMa* package (*60*). All models reasonably fit the data when looking at residuals and Q-Q plots. Although there were sometimes slight deviations from model assumptions, transformations did not improve fit, and mixed effects models are generally robust to slight deviations (*61*).

Next, we wanted to test how ovarian fluid and temperature interact to influence sperm longevity of the different male types. Longevity could also be important in terms of fertilization success not necessarily for the initial clutch of eggs, but future clutches of eggs due to rapid sequential matings of the same or different females in this system. To assess longevity, we looked at sperm velocity at five minutes. We chose this time to be consistent with previous work in this system (*26*). We also did not do a time series of sperm velocity to avoid the complexity of interpreting and analyzing a potential 4-way interaction among time, activating fluid, experimental temperature, and male type. For this analysis, we had 12 complete nesting male replicates and 14 complete sneaker male replicates. We had measurements for four or five treatments for the incomplete replicates. We included the incomplete replicates because mixed effects models account for incomplete blocks, and the data can be used to estimate the treatment effects they contained (see Table S2 for a summary of sample size by treatment). The average number of sperm used was 91.5± 6.24 per measurement. We ran the same statistical analyses as the initial sperm velocity analysis.

Because temperature changes throughout the reproductive season (Fig. 1), fish may be acclimated to different temperatures when used in the experiment. We thus ran supplementary analyses that included a fixed effect of the difference in ocean water temperature on the day of the experiment and trial. This effect did not influence any of the results (Table S9), did not improve model fit (Table S10), and was highly correlated with temperature treatment (*r* = 0.9775). Therefore, we did not include this effect in any of the results discussed in the main text. We conducted all analyses and made all figures in R V.4.1.2 (*56*) using the “tidyverse” suite of packages (*62*).

### Statistical analysis of variation

Understanding individual differences in responses is important to understand the evolutionary potential of the population. We tested if there was significant variation among males in the response of sperm to either ovarian fluid or temperature. To do this, we used a log-likelihood ratio test comparing models with only random intercepts to models with a random slope for activating fluid treatment and water test temperature. We performed this analysis for both sperm velocity at 30 seconds and at five minutes.

To estimate differences in variation between temperature treatments, we generated a posterior distribution of 10,000 random draws of the standard deviation of the random slopes using the *arm* package (*63*). To ease interpretation, we subset the data by activating fluid treatment and male type before refitting the models and dropping the random intercept term. For each model, we extracted the standard deviation of the random slope of each temperature treatment. In each posterior draw, we calculated the difference in standard deviation to generate a posterior distribution of the differences in standard deviation at different temperature treatments.

### Estimated paternity

To determine how temperature and activating fluid influence the relative paternity of nesting males, we used data from (*26*) with a generalized linear model with a quasibinomial family to account for overdispersion. We modeled nesting male paternity as a function of relative sperm velocity, relative sperm number, fluid (seawater or ovarian fluid), and the interaction between fluid and relative sperm number. For relative sperm numbers, we used the average sperm number released by the different male types in natural matings from (*45*). For relative sperm velocity, we used the sperm velocity estimates at the different temperatures and activating fluids. We accounted for error in our sperm velocity estimates at different temperatures and activating fluids and error in the paternity statistical model by generating a posterior distribution with the *arm* package with 10,000 random draws (*63*).

### Supplemental Tables

**Table S1.** Sample sizes for sperm velocity analysis at 30 seconds.

| Male tactic | Activating fluid | Test temperature | Sample size |
| --- | --- | --- | --- |
| Nesting Male | Seawater | 16°C | 17 |
| Nesting Male | Seawater | 22°C | 18 |
| Nesting Male | Seawater | 28°C | 19 |
| Nesting Male | Ovarian fluid | 16°C | 18 |
| Nesting Male | Ovarian fluid | 22°C | 18 |
| Nesting Male | Ovarian fluid | 28°C | 19 |
| Sneaker | Seawater | 16°C | 18 |
| Sneaker | Seawater | 22°C | 19 |
| Sneaker | Seawater | 28°C | 19 |
| Sneaker | Ovarian fluid | 16°C | 18 |
| Sneaker | Ovarian fluid | 22°C | 19 |
| Sneaker | Ovarian fluid | 28°C | 18 |

**Table S2.**
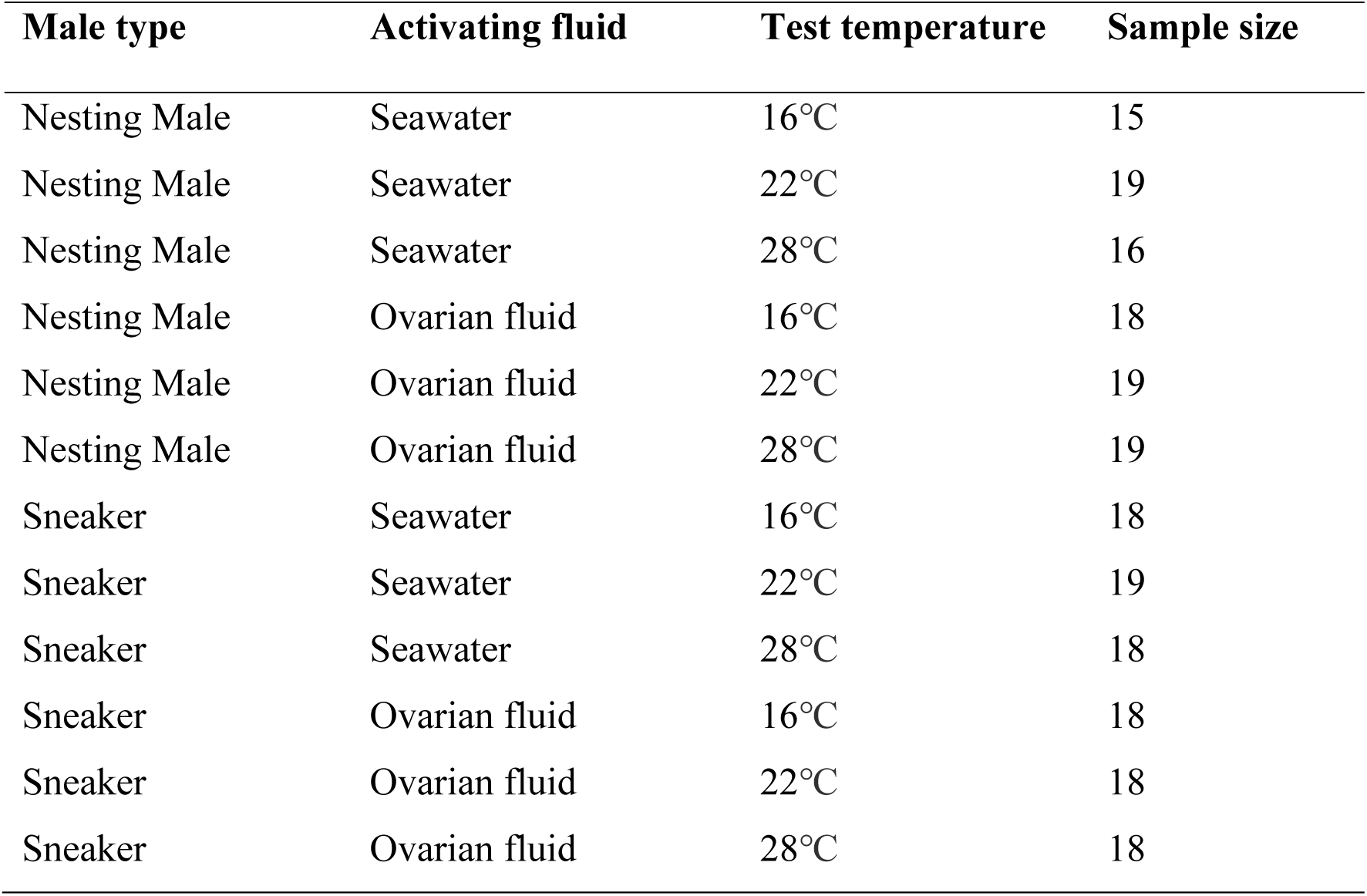
Sample sizes for sperm velocity analysis at five minutes.

**Table S3.** Temperature contrasts for sperm velocity at 30 seconds. Bolded rows are significant.

| Contrast | Activating fluid | Male | Estimate | SE | Z ratio | *p*(\|Z\|>0) |
| --- | --- | --- | --- | --- | --- | --- |
| **16 - 22°C** | **Seawater** | **Nesting Male** | **-35.573** | **8.032** | **-4.429** | **<0.001** |
| **16 - 28°C** | **Seawater** | **Nesting Male** | **-50.217** | **8.926** | **-5.626** | **<0.001** |
| 22 - 28°C | Seawater | Nesting Male | -14.645 | 8.349 | -1.754 | 0.185 |
| 16 - 22°C | Seawater | Sneaker | -16.875 | 7.855 | -2.148 | 0.08 |
| **16 - 28°C** | **Seawater** | **Sneaker** | **-58.75** | **8.636** | **-6.803** | **<0.001** |
| **22 - 28°C** | **Seawater** | **Sneaker** | **-41.875** | **8.526** | **-4.911** | **<0.001** |
| 16 - 22°C | Ovarian fluid | Nesting Male | 8.62 | 7.968 | 1.082 | 0.525 |
| 16 - 28°C | Ovarian fluid | Nesting Male | -8.847 | 8.749 | -1.011 | 0.570 |
| 22 - 28°C | Ovarian fluid | Nesting Male | -17.467 | 8.721 | -2.003 | 0.112 |
| **16 - 22°C** | **Ovarian fluid** | **Sneaker** | **-23.503** | **7.818** | **-3.006** | **0.007** |
| **16 - 28°C** | **Ovarian fluid** | **Sneaker** | **-38.184** | **8.763** | **-4.357** | **<0.001** |
| 22 - 28°C | Ovarian fluid | Sneaker | -14.682 | 8.664 | -1.695 | 0.207 |

**Table S4.** Activating medium contrasts for sperm velocity at 30 seconds. Bolded rows are significant.

| Contrast | Temperature | Male | Estimate | SE | Z ratio | $p(\vert Z\vert >0)$ |
| --- | --- | --- | --- | --- | --- | --- |
| **OF - SW** | **16°C** | **Nesting Male** | **37.548** | **8.01** | **4.688** | **<0.001** |
| OF - SW | 22°C | Nesting Male | -6.645 | 7.918 | -0.839 | 0.401 |
| OF - SW | 28°C | Nesting Male | -3.823 | 7.812 | -0.489 | 0.625 |
| **OF - SW** | **16°C** | **Sneaker** | **15.719** | **7.809** | **2.013** | **0.044** |
| **OF - SW** | **22°C** | **Sneaker** | **22.347** | **7.575** | **2.95** | **0.003** |
| OF - SW | 28°C | Sneaker | -4.846 | 7.778 | -0.623 | 0.533 |

**Table S5.** Male type contrasts for sperm velocity at 30 seconds. Bolded rows are significant.

| Contrast | Temperature | Activating medium | Estimate | SE | Z ratio | $p(\vert Z\vert >0)$ |
| --- | --- | --- | --- | --- | --- | --- |
| NM - SN | 16°C | Seawater | -11.406 | 9.307 | -1.226 | 0.22 |
| NM - SN | 22°C | Seawater | 7.291 | 9.821 | 0.742 | 0.458 |
| NM - SN | 28°C | Seawater | -19.939 | 13.732 | -1.452 | 0.146 |
| NM - SN | 16°C | Ovarian fluid | 10.423 | 8.261 | 1.262 | 0.207 |
| **NM - SN** | **22°C** | **Ovarian fluid** | **-21.701** | **9.067** | **-2.393** | **0.017** |
| NM - SN | 28°C | Ovarian fluid | -18.915 | 9.847 | -1.921 | 0.055 |

**Table S6.**
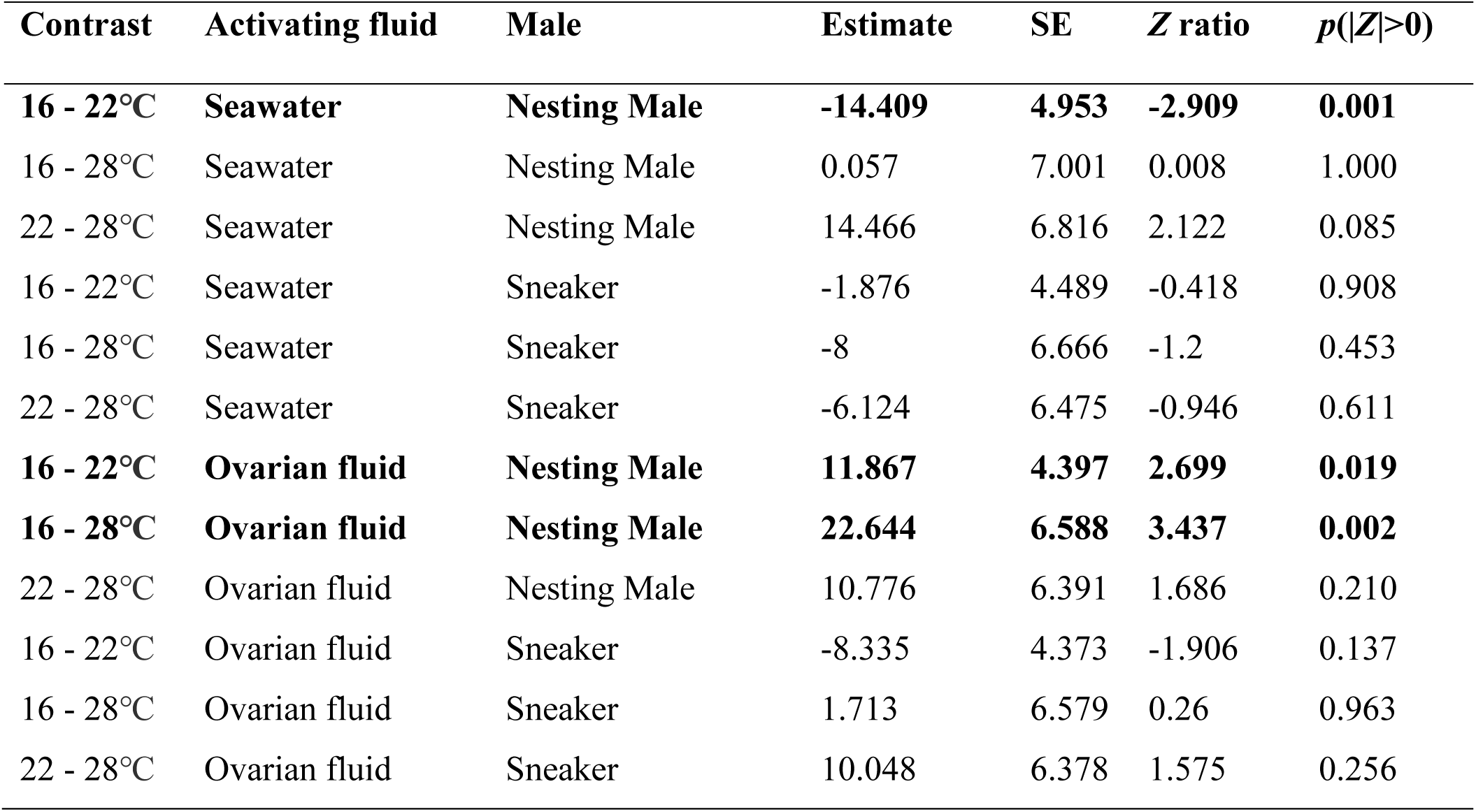
Temperature contrasts for sperm velocity at five minutes. Bolded rows are significant.

**Table S7.** Activating medium contrasts for sperm velocity at five minutes. Bolded rows are significant.

| Contrast | Temperature | Male | Estimate | SE | Z ratio | $p(\vert Z\vert >0)$ |
| --- | --- | --- | --- | --- | --- | --- |
| **OF - SW** | **16°C** | **Nesting Male** | **41.767** | **6.417** | **6.508** | **<0.001** |
| **OF - SW** | **22°C** | **Nesting Male** | **15.491** | **6.378** | **2.429** | **0.015** |
| **OF - SW** | **28°C** | **Nesting Male** | **19.18** | **6.485** | **2.958** | **0.003** |
| **OF - SW** | **16°C** | **Sneaker** | **18.099** | **6.185** | **2.926** | **0.003** |
| **OF - SW** | **22°C** | **Sneaker** | **24.558** | **6.209** | **3.955** | **<0.001** |
| OF - SW | 28°C | Sneaker | 8.386 | 6.267 | 1.338 | 0.181 |

**Table S8.**
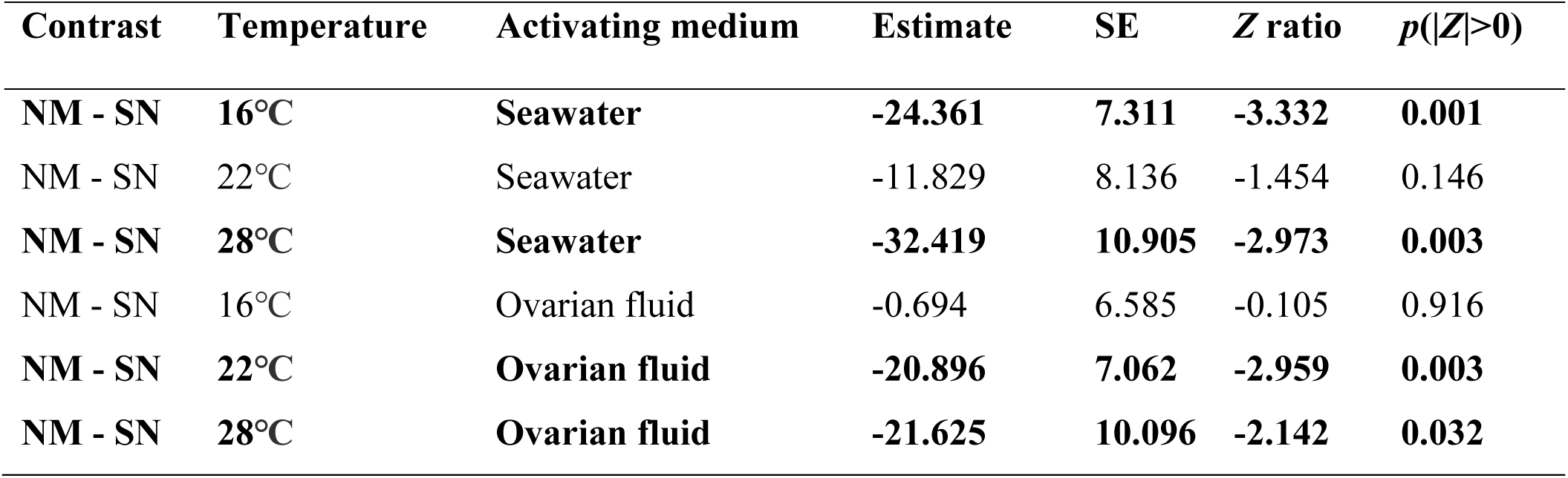
Male type contrasts for sperm velocity at five minutes. Bolded rows are significant.

| Contrast | Temperature | Activating medium | Estimate | SE | Z ratio | $p(\vert Z\vert >0)$ |
| --- | --- | --- | --- | --- | --- | --- |
| NM - SN | 16°C | Seawater | -24.361 | 7.311 | -3.332 | 0.001 |
| NM - SN | 22°C | Seawater | -11.829 | 8.136 | -1.454 | 0.146 |
| NM - SN | 28°C | Seawater | -32.419 | 10.905 | -2.973 | 0.003 |
| NM - SN | 16°C | Ovarian fluid | -0.694 | 6.585 | -0.105 | 0.916 |
| NM - SN | 22°C | Ovarian fluid | -20.896 | 7.062 | -2.959 | 0.003 |
| NM - SN | 28°C | Ovarian fluid | -21.625 | 10.096 | -2.142 | 0.032 |

**Table S9.** Results of linear mixed-effects models explaining sperm velocity at (A) 30 seconds and (B) five minutes including the effect of the difference between water temperature in which the fish were caught and test temperature. The significance of fixed effects was determined with a Type II ANOVA Wald χ^2^ test. Random effects for both models were a random intercept for the field of view of the video nested within the trial and random slopes for both activating fluid treatment and temperature treatment of each trial. Significant effects are bolded.

| Trait | Model Effect | $\chi^2$ | $p(>\chi^2)$ |
| --- | --- | --- | --- |
| (A) Sperm velocity at 30 seconds | **Temperature** | **13.8911** | **0.0010** |
|  | Activating fluid | 3.6865 | 0.0549 |
|  | **Difference in water temperature** | **4.2856** | **0.0384** |
|  | Male type | 0.3777 | 0.5388 |
|  | **Activating fluid: test temperature** | **61.1165** | **<0.0001** |
|  | Activating fluid: male type | 0.0877 | 0.7671 |
|  | Test temperature: male type | 3.4845 | 0.1751 |
|  | **Activating fluid: test temperature: male type** | **41.9645** | **<0.0001** |
| (B) Sperm velocity at five minutes | Temperature | 1.8608 | 0.3944 |
|  | **Activating fluid** | **24.6223** | **<0.0001** |
|  | Difference in water temperature | 0.6418 | 0.4231 |
|  | **Male type** | **6.0445** | **0.0140** |
|  | **Activating fluid: test temperature** | **53.2031** | **<0.0001** |
|  | Activating fluid: male type | 0.8464 | 0.0876 |
|  | Test temperature: male type | 4.8710 | 0.0730 |
|  | **Activating fluid: test temperature: male type** | **74.6250** | **<0.0001** |

**Table S10.** Including the difference in water temperature between which the fish were caught and the test temperature did not significantly improve model fit. Results of the log-likelihood ratio tests of sperm velocity at (**A**) 30 seconds and (**B**) five minutes.

| Trait | Model | AIC | $\chi^2$ | P(>$\chi^2$) |
| --- | --- | --- | --- | --- |
| (A) Sperm velocity 30 seconds | Base model | 524555 | NA | NA |
|  | With temp diff | 524553 | 3.7568 | 0.0526 |
| (B) Sperm velocity five minutes | Base model | 198266 | NA | NA |
|  | With temp diff | 198267 | 0.668 | 0.4137 |

**Table S11.**
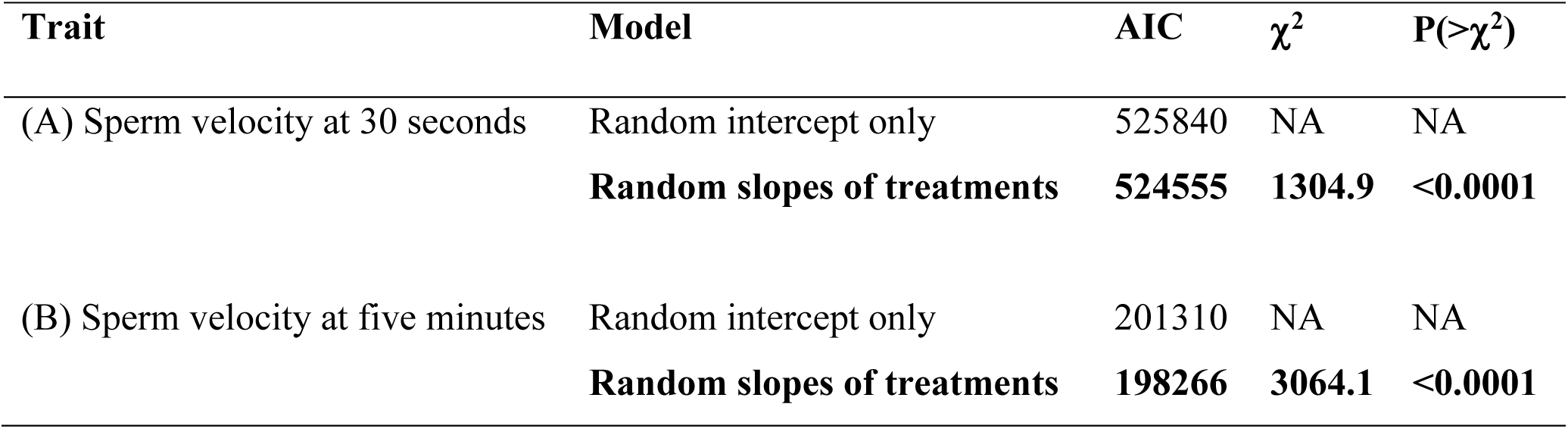
Results of the log-likelihood ratio tests of sperm velocity at (A) 30 seconds and (B) five minutes. Models that significantly improved fit are bolded.

### Supplemental Figures

**Fig. S1.**
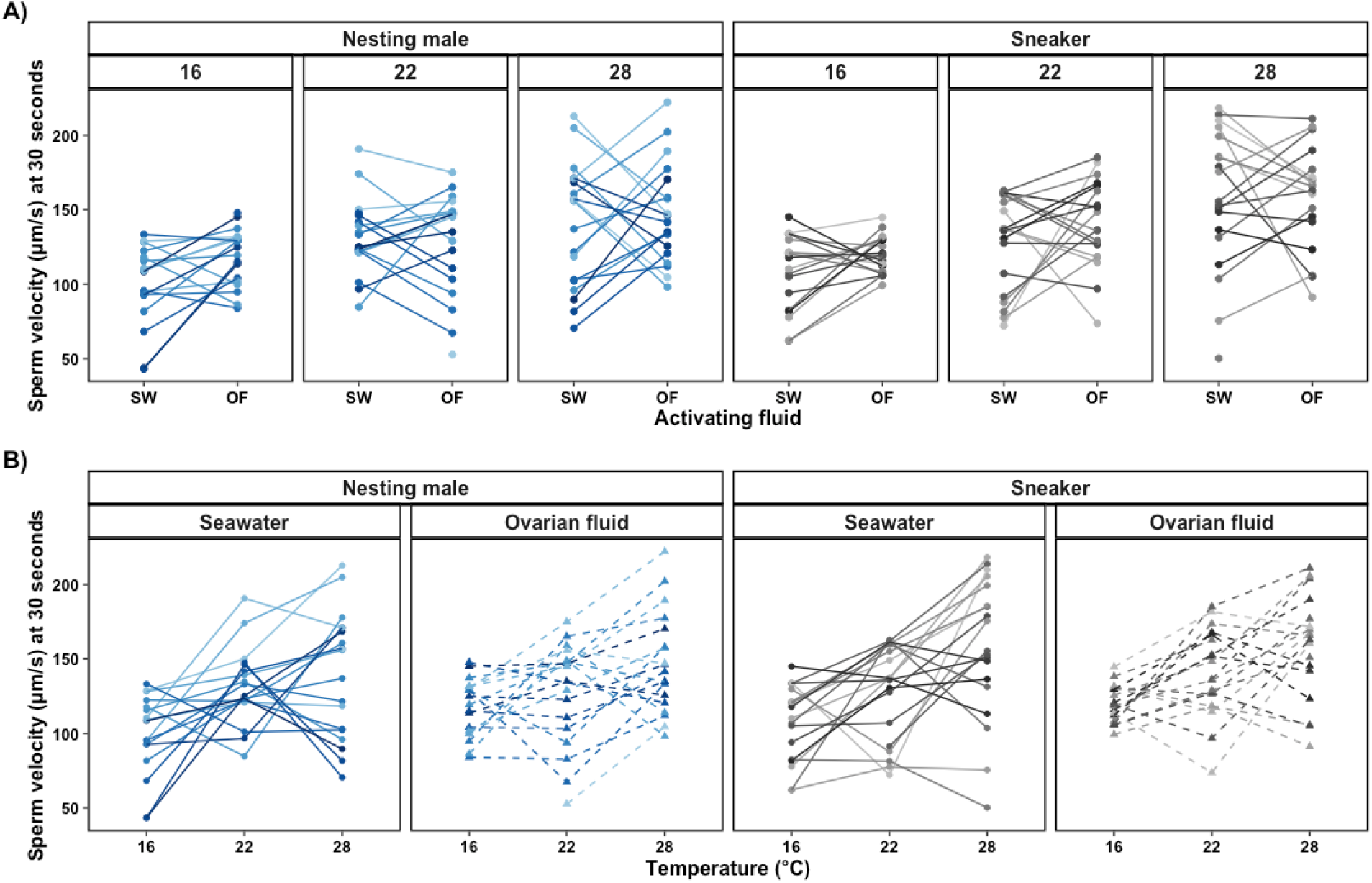
There is significant variation in the effects of ovarian fluid and temperature across experimental replicates on sperm velocity at 30 seconds. Each line of connected points and color represents a single replicate of the experiment (the same male and female pair). Visualization of reaction norms for **(A)** activating fluid and **(B)** temperature. Both A and B have the same data. For this visualization, we excluded incomplete replicates. The points are the average sperm velocity at 30 seconds.

**Fig. S2.**
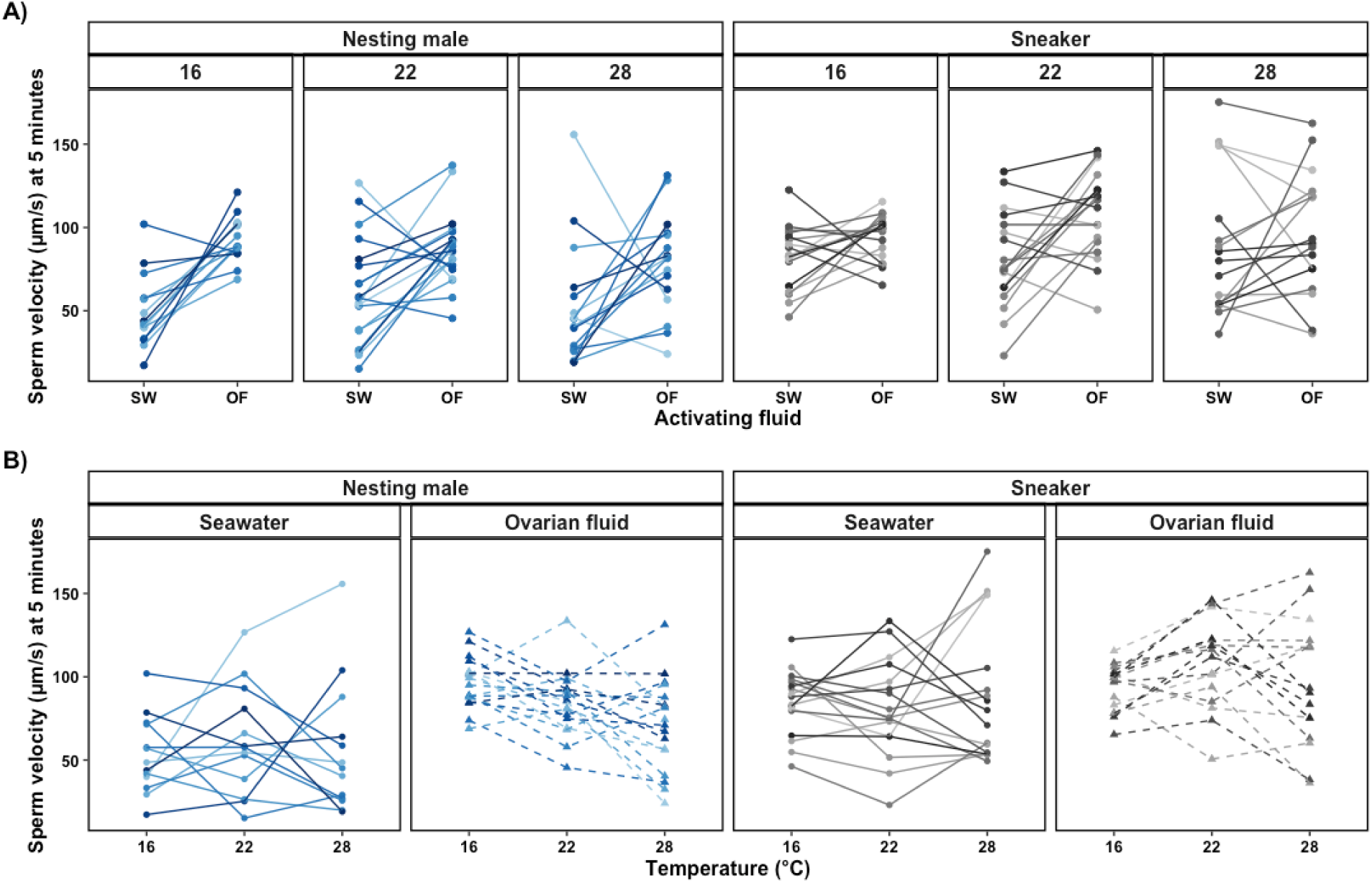
There is significant variation in the effects of ovarian fluid and temperature across experimental replicates on sperm velocity at five minutes. Each line of connected points and color represents a single replicate of the experiment (the same male and female pair). Visualization of reaction norms for **(A)** activating fluid and **(B)** temperature. Both A and B have the same data. For this visualization, we excluded incomplete replicates. The points are the average sperm velocity at five minutes.

**Fig. S3.**
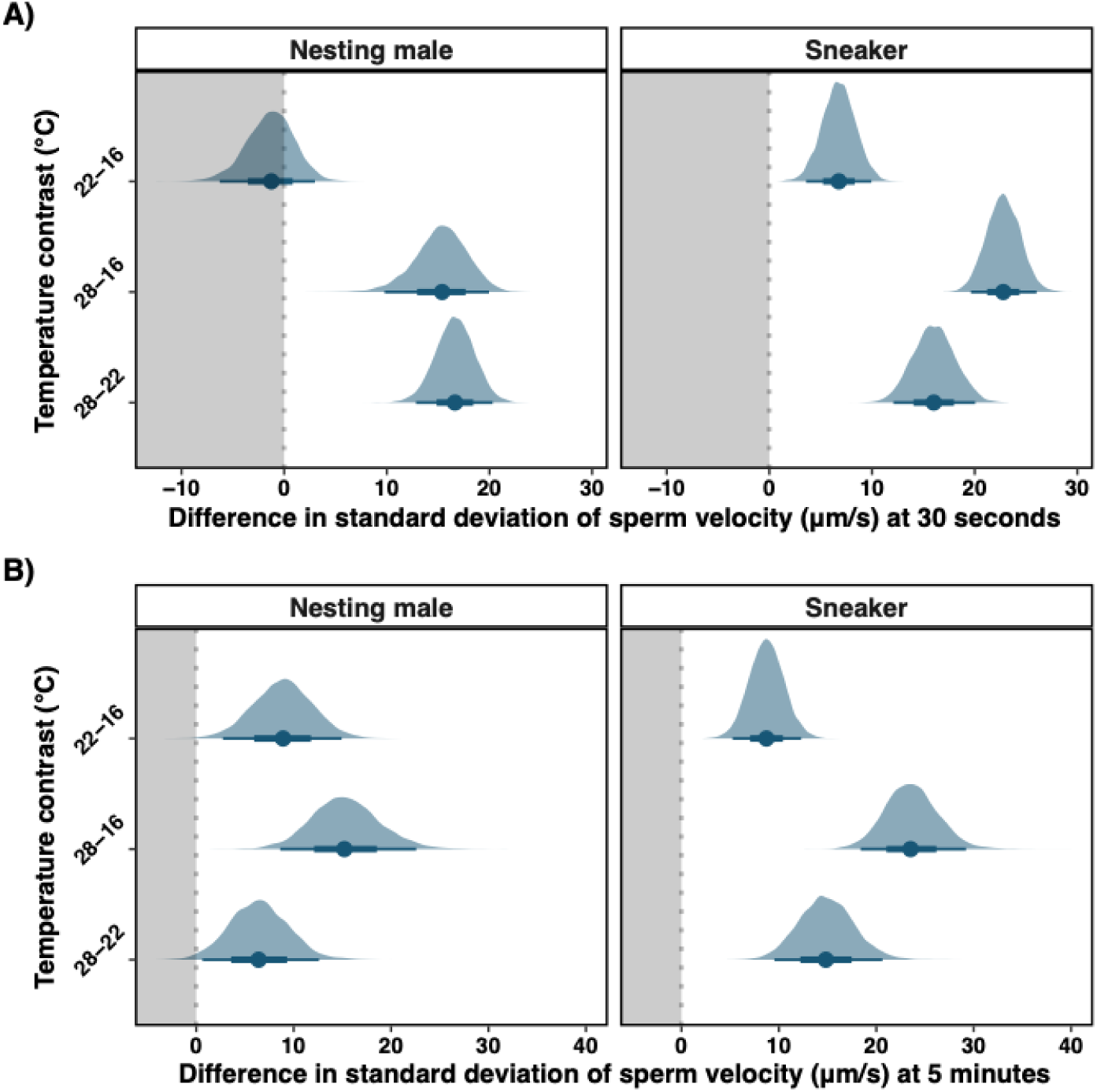
Variation in sperm velocity generally increases with temperature in seawater. Posterior probability distributions of the difference in random slope standard deviation at different temperatures for nesting males (left) and sneakers (right) in seawater at **(A)** 30 seconds and **(B)** five minutes. The posterior distributions were generated with 10,000 simulated draws. Solid points represent the median of the posterior, thin bars represent the 95% credible intervals of the posterior distribution, and thick bars represent the 66% credible interval. The dotted grey lines at zero mean no difference in standard deviation. Negative values (grey background) indicate higher temperature decreased variation. Positive values (white background) indicate higher temperatures increased variation. Similar plots for ovarian fluid treatment are shown in Fig. 3.

**Fig. S4.**
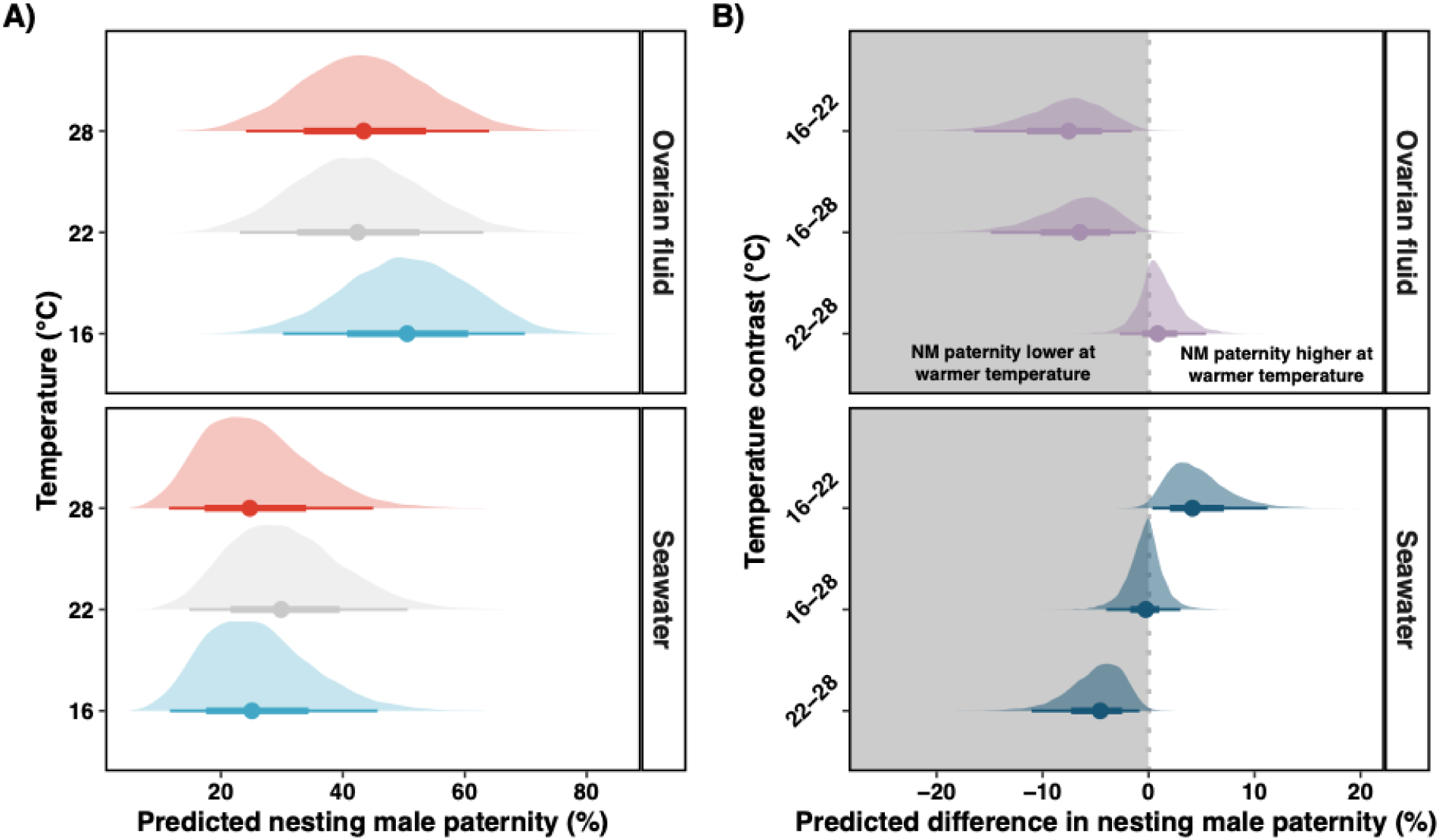
Ovarian fluid increases nesting male paternity, but this effect is weakened at warmer temperatures. **(A)** Shown are the posterior probability distributions of predicted nesting male paternity at 16℃, 22℃, and 28℃ in either ovarian fluid (top) or seawater (bottom). **(B)** Ovarian fluid has a less positive impact on nesting male (NM) paternity at 22℃ and 28℃. The posterior probability distributions of the difference in nesting male paternity at different temperatures in either ovarian fluid (top) or seawater (bottom) are shown. For all panels, solid points represent the median of the posterior, thin bars represent the 95% credible intervals of the posterior distribution, and thick bars represent the 66% credible interval. The posterior distributions were generated with 10,000 simulated draws. The dotted grey lines at zero mean no difference in nesting male paternity. Negative values (grey background) indicate higher temperature decreased nesting male paternity. Positive values (white background) indicate higher temperatures increased nesting male paternity. The top subpanel of B is the same as Fig. 4 in the main text.

